# Intrinsic and extrinsic noise are distinguishable in a synthesis – export – degradation model of mRNA production

**DOI:** 10.1101/2020.09.25.312868

**Authors:** Gennady Gorin, Lior Pachter

## Abstract

Intrinsic and extrinsic noise sources in gene expression, originating respectively from transcriptional stochasticity and from differences between cells, complicate the determination of transcriptional models. In particularly degenerate cases, the two noise sources are altogether impossible to distinguish. However, the incorporation of downstream processing, such as the mRNA splicing and export implicated in gene expression buffering, recovers the ability to identify the relevant source of noise. We report analytical copy-number distributions, discuss the noise sources’ qualitative effects on lower moments, and provide simulation routines for both models.

## 1 Background

Recent improvements in single-cell transcriptomics, including increasingly sensitive fluorescence- and sequencing-based methods, have begun to provide data useful for discriminating between competing biophysical models. One immediate application of interest is that of *intrinsic* and *extrinsic* cellular gene expression noise, which has already been studied directly from mRNA reporter statistics [1, 2]. While experimental and statistical methods for measuring the relative contributions of intrinsic and extrinsic noise are relatively advanced [3–6], microscopic models of cell-to-cell variability are less well developed. These models are necessary in light of recent methods for measuring the molecular state of cells, which offer routes to better mechanistic understanding, but present a number of new challenges in controlling noise sources.

While the introduction of single-cell RNA sequencing (scRNA-seq) data with unique molecular identifiers (UMIs) provides measurements of a substantial fraction of transcripts in individual cells [7], the resulting copy-number data are discrete, and thus challenging to model with existing methods that largely focus on continuous-valued fluorescence readouts. The biochemistry of scRNA-seq also generally relies on the capture of polyadenylated sequences in fixed media [8], which limits the scope of assays, and is not directly compatible with *in vivo* experimental methods relying on the integration of multiple fluorescent reporters to distinguish between the sources of noise [3]. Furthermore, the analysis of lower moments of gene expression has been shown to be insufficient for the identification of biophysical parameters even for purely intrinsic noise models [9], suggesting that full copy-number distributions are necessary for modeling more complex systems with multiple sources of noise.

Another challenge lies in theory; ideally, analytical results will be available to provide qualitative interpretability and guide computational approaches, but many current methods are purely numerical. For example, while methods for the explicit description of extrinsic noise are formally available, in the context of a transcriptional model, the incorporation of extrinsic noise typically corresponds to the construction of a mixture model with parameter values drawn from a distribution [2, 3, 10]. Under this construction, full analytical solutions are only available in the simplest cases.

## 2 Two models for gene expression

It is well-known that the common two-state model of gene expression [11] gives rise to a negative binomial (NB) distribution of mRNA counts in the short-burst limit [12]. However, a recent study shows that constitutive transcription in a cell population with a gamma-distributed production rate parameter also yields a negative binomial distribution of mRNA counts [13]. Although there are both experimental and theoretical arguments favoring a bursting model for eukaryotic transcription [14–17] – current theories posit that superstructure modifications are responsible for occlusion and exposure of the gene locus [18, 19] – a comprehensive model should account for all relevant sources of noise, as well as provide both a quantitative and qualitative understanding of their effects.

A current limitation of existing models is that processes downstream of eukaryotic mRNA production, such as export and/or splicing processes [20, 21], are generally ignored. However, promising new technologies and experiments, based on fluorescence [22, 23] and sequencing [24] methods, can distinguish nascent from mature mRNA molecules based on spatial or intronic readouts, thus providing essential data for studying model of increasing complexity. In particular, the maturation of these methods and the availability of resulting multimodal data naturally suggests the potential of fitting otherwise poorly identifiable models [9].

As a first step, and to gain a qualitative understanding of the effects of intrinsic and extrinsic noise in the context of downstream processing, we compare solutions of two simple two-stage models of transcription that include downstream processing at steady state. Both models assume that nascent mRNA (unspliced or pre-mRNA) is converted to mature mRNA (spliced mRNA) after an exponentially-distributed delay, corresponding to splicing. This is followed by another exponentially-distributed delay that models the mature mRNA being degraded. The splicing rate *β* and degradation rate *γ* are deterministic. The gene locus dynamics are modeled by either bursts, with stochastic burst size *B* ∼ *Geom*(*b*) and deterministic burst *i*nitiation frequency *k*_*i*_ [20], or constitutive, with stochastic but constant transcription rate *K* ∼ *Gamma*(*α, η*). The model parametrizations are illustrated in Figure 1. We calculate lower moments and cross-moments, and show how these can be used to differentiate between distributions and statistics resulting from the two models.

**Figure 1:**
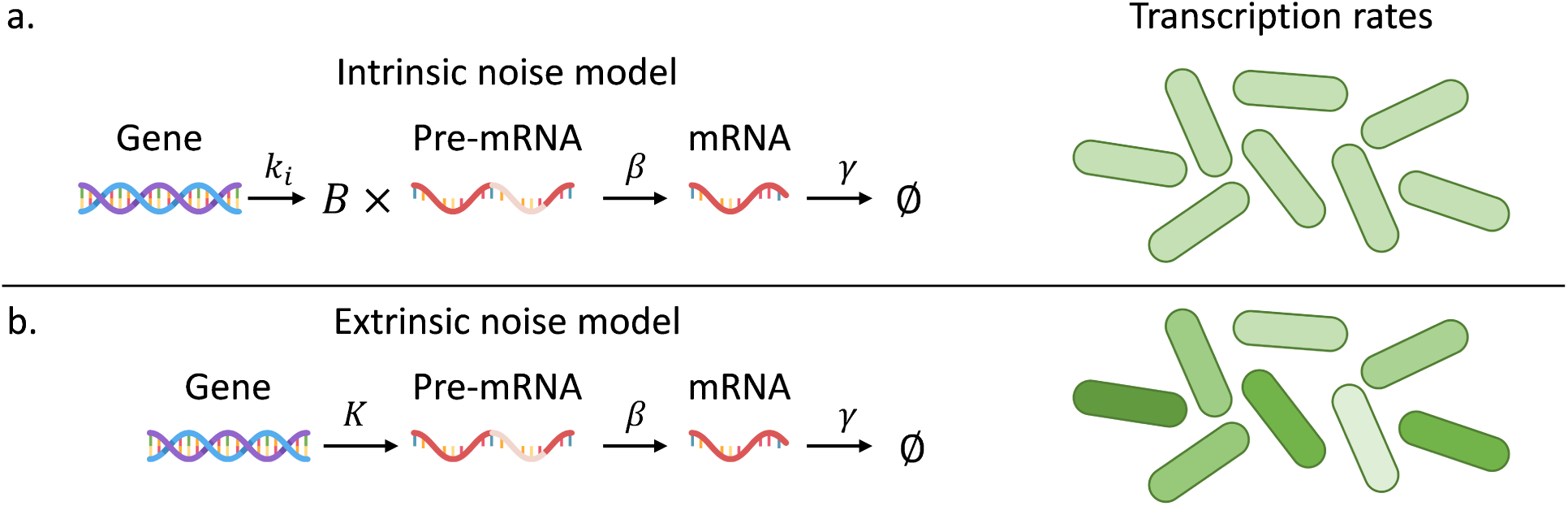
(a) Schema of the intrinsic noise model (*k*_*i*_: burst frequency; *B*: burst size drawn from a geometric distribution; *β*: pre-mRNA splicing rate; *γ*: mRNA degradation rate. Uniform shade of green indicates identical parameter values across all cells). (b) Schema of the extrinsic noise model (*K*: transcription rate; *β*: pre-mRNA splicing rate; *γ*: mRNA degradation rate. Different shades of green indicate different values of *K* across cells).

## 3 Notation

### 3.1 Model parametrization

Model-independent quantities and statistics are defined in Table 1. The two models’ parameters are defined in Tables 2 and 3. Finally, *x*_*z*_, where *x* is a statistic computed from data moments (e.g., *µ*_*M*_, 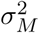, *ρ, γ*) and *z ∈* {*i, e*} refers to the predicted value of that statistic based on either the *i*ntrinsic or *e*xtrinsic noise model. For example, *µ*_*M,i*_ refers to the predicted mean mature mRNA copy number under the intrinsic noise model, while *ρ*_*e*_ refers to the predicted nascent–mature correlation under the extrinsic noise model.

**Table 1:** Observation variables

| Parameter | Definition |
| --- | --- |
| $n$ | Number of nascent mRNA |
| $m$ | Number of mature mRNA |
| $N$ | Random variable denoting number of nascent mRNA; $N \in \mathbb{N}_0$ |
| $M$ | Random variable denoting number of mature mRNA; $M \in \mathbb{N}_0$ |
| $\gamma$ | Degradation rate |
| $\mu_z, \mathbb{E}[Z]$ | Expectation of species $Z \in \{N, M\}$ |
| $\mu$ | Expectation of an arbitrary distribution |
| $\sigma_z^2$ | Variance of species $Z \in \{N, M\}$ |
| $\sigma^2$ | Variance of an arbitrary distribution $Z \in \{N, M\}$ |
| $Cov(N, M)$ | Covariance nascent and mature copy numbers |
| $\rho$ | Pearson correlation coefficient |
| $q$ | Computed statistic $\frac{\sigma_N^2}{\mu_N} - 1$ |
| $P(n, m; \cdot)$ | Joint PMF of nascent and mature mRNA, shorthand for $P(N = n, M = m; \cdot)$ |
| $P(\cdot; \cdot)$ | PMF of an arbitrary parametrized discrete random variable |
| $f(\cdot; \cdot)$ | PDF of an arbitrary parametrized continuous random variable |

**Table 2:** Intrinsic noise model parameters

| Parameter | Definition |
| --- | --- |
| $k_i$ | Burst frequency |
| $B$ | Geometric random variable denoting burst size |
| $b$ | Mean of $B$ |
| $\beta$ | Splicing rate |
| $\gamma$ | Degradation rate |
| $f$ | $\frac{\beta}{\beta+\gamma}$, non-dimensional splicing rate |

**Table 3:** Extrinsic noise model parameters

| Parameter | Definition |
| --- | --- |
| $K$ | Gamma-distributed random variable denoting transcription rate |
| $\alpha$ | Shape of gamma distribution |
| $\eta$ | Rate of gamma distribution |
| $\beta$ | Splicing rate |
| $\gamma$ | Degradation rate |
| $f$ | $\frac{\beta}{\beta+\gamma}$, non-dimensional splicing rate |

A probability mass function (PMF) associated with a discrete-valued random variable *X* is equivalently denoted by *P* (·; ·) or *P* (*X* = *k*; ·). A probability density function (PDF) associated with a continuous-valued random variable is denoted by *f* (·; ·).

### 3.2 Probability distributions

The geometric distribution is defined as follows: if *X* ∼ *Geom*(*p*), *P* (*X* = *k*; *p*) = (1 *− p*)^*k*^*p*, where *p ∈* (0, 1] and *k ∈* ℕ_0_. The geometric distribution is well-known to arise in the short-burst limit of the two-state transcription model [25].

The negative binomial distribution is defined as follows: if *X* ∼ *NegBin*(*r, p*), 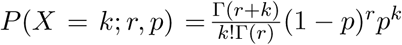, where *p ∈* [0, 1] and *r >* 0. We note that MATLAB and the NumPy library take the opposite convention, with a 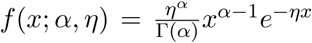 parameter defined as 1 *− p*.

The gamma distribution is defined as follows: if *X* ∼ *Gamma*(*α, η*), 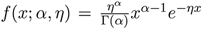.

This is the shape/rate parametrization. We note that MATLAB and the NumPy library take the opposite shape/scale parametrization with parameter *θ* = *η*^*−*1^. Furthermore, the rate *η* is usually given the variable name “*β*”; however, we use the current convention to preclude confusion with the splicing rate parameter.

## 4 Preliminaries

### 4.1 Intrinsic noise model

#### 4.1.1 Probability mass function

The full joint distribution for the burst model requires numerical integration and Fourier transformation [20]. To our knowledge, no analytical solution exists, although approximations in terms of hypergeometric functions are available [26].

The nascent marginal is distributed per 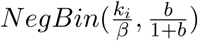. The mature marginal is NB-distributed in the limit of low *β* and Poisson-distributed in the limit of high *β*. Although the distribution in the intermediate region is qualitatively similar to NB, it does not appear to be *exactly* representable as NB. Furthermore, even the determination of the closest NB approximation according to some divergence metric is an open problem, although method of moments approximations may be satisfactory for some purposes.

#### 4.1.2 Moments

Per the results from Singh and Bokes [20]:

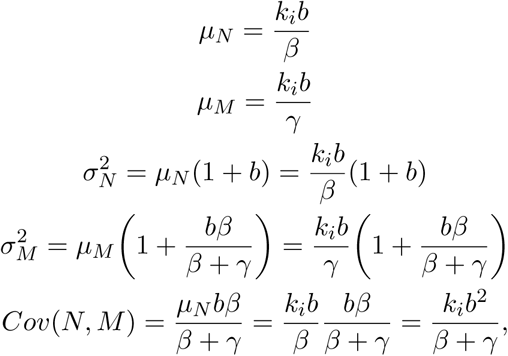

yielding the following Pearson correlation coefficient:

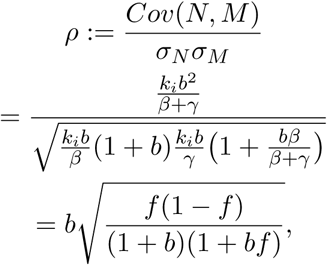

where *f* controls the relationship between the splicing and degradation timescales.

### 4.2 Extrinsic noise model

#### 4.2.1 Probability mass function

The full time-dependent copy-number probability distribution under constitutive production is well-known and represents one of the most valuable and general results in chemical master equation (CME) analysis [27]. In the relevant steady-state regime, the solution 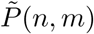 giving the probability of a state with a given number of nascent and mature molecules is the product of independent Poisson distributions. Given a production rate *K*, splicing rate *β*, and degradation rate *γ*,

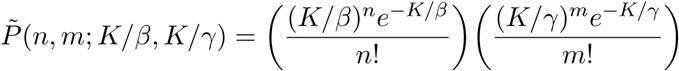

Therefore, marginalizing over the gamma-distributed production rate *K*:

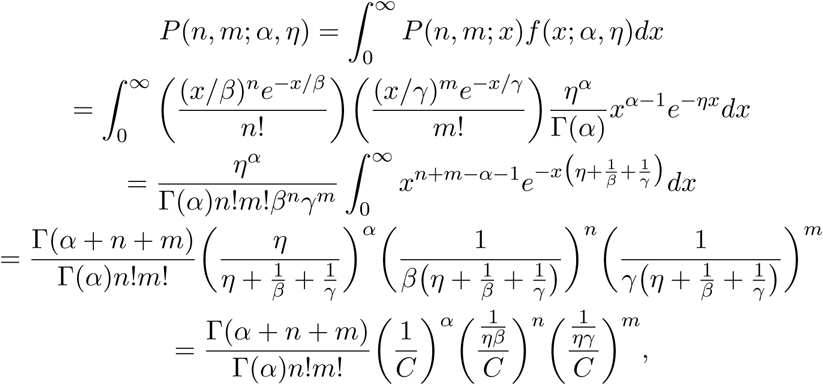

where 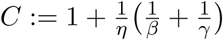. This is the multivariate negative binomial (MVNB) distribution [28]. For the sake of completeness, we show that the marginal distributions take the expected negative binomial form:

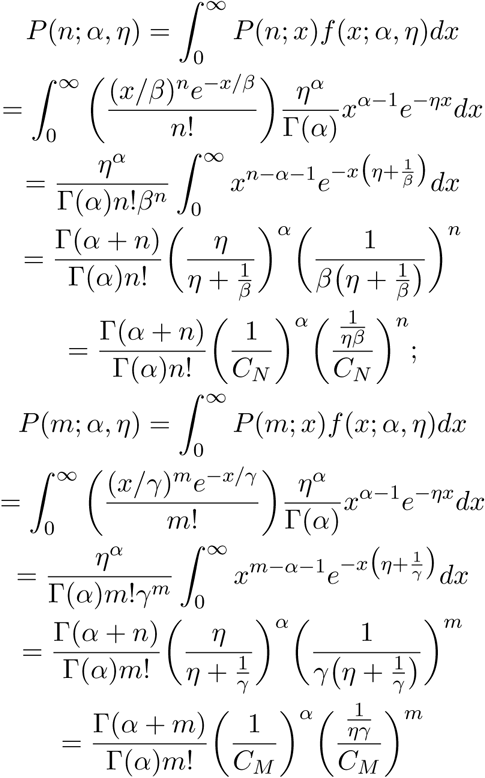

where 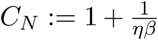 and 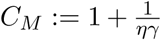. The two marginals’ NB parameters are 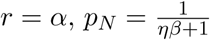 and 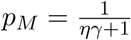.

We note that the Poissonian framework due to Jahnke and Huisinga [27] yields the solutions for arbitrary graphs representing sources, sinks, and reaction channels. This is sufficient, for example, to construct a directed acyclic graph representing alternative splicing of a constitutively expressed gene. Adding extrinsic noise to these graphs is trivial and immediately follows from the definitions of the corresponding Poisson rate constants.

#### 4.2.2 Moments

The moments and variances of the marginals follow immediately from standard identities for the NB distribution:

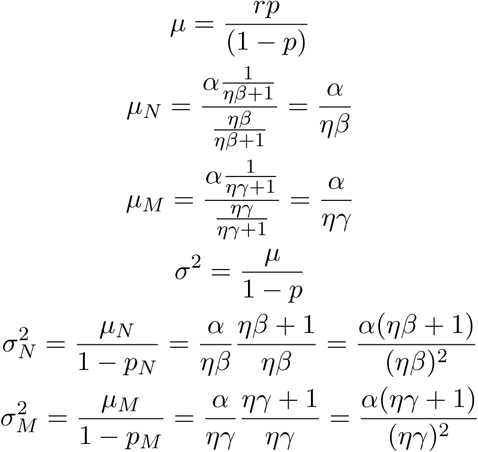

The moment-generating function (MGF) of the MVNB distribution is 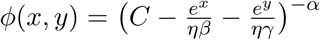 [28]. Differentiating the expression with respect to *x* and *y* yields 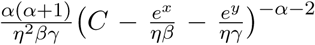. Evaluating at *x* = *y* = 1 yields the cross moment 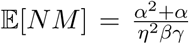. Therefore, the covariance is 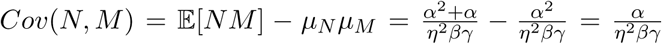. This result yields the following Pearson correlation coefficient:

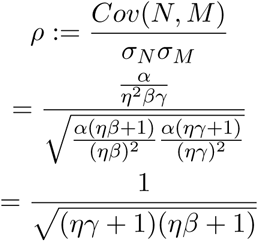

## 5 Discriminating between intrinsic and extrinsic noise models

Using the above computations, we can show that steady-state information about the nascent and mature distributions is sufficient to distinguish between the two models. We start from an *a priori* non-identifiable negative binomial nascent mRNA distribution, and demonstrate disagreement between statistics predicted for the mature mRNA. For the purposes of illustration, we assume data-based constraints upon *µ*_*M*_ and upon 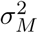, motivated by the existence of experimental methods for determining these quantities [1, 29]. However, we restrict our analysis to analytical distributions to avoid the details of any particular observation or statistical inference method.

The intrinsic and extrinsic noise models are respectively parametrized by {*b, k*_*i*_, *β, γ*} and {*α, η, β, γ*}. However, at steady state, the time variable is not independently identifiable. Therefore, the absolute scaling of the rate variables {*k*_*i*_, *η, β, γ*} is not feasible to determine. This non-identifiability is self-evident from the functional forms of the distributions, e.g. the moment dependence on 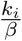 and 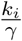 in the intrinsic noise model and on *ηβ* and *ηγ* in the extrinsic noise model. Therefore, we set *β* to 1 with no loss of generality.

Further, the nascent marginal is governed by the two-parameter NB distribution. In the context of model fitting, this implies that two of the parameters of the joint distribution are fully determined by the nascent distribution, and only one degree of freedom remains to be determined by the mature mRNA data.

Crucially, given a negative binomial distribution of nascent mRNA, with mean *µ*_*N*_ and variance 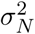, the intrinsic and extrinsic noise models are not distinguishable. Using the intrinsic noise model uniquely identifies 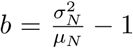 and 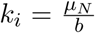. Conversely, using the extrinsic noise model uniquely identifies 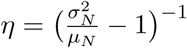 and *α* = *µ*_*N*_*η*.

### 5.1 Case of constrained *γ* or *µ*_*M*_

Constraining *γ* is equivalent to fixing the mature mRNA means:

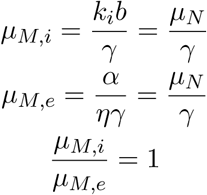

However, the higher moments disagree. Defining the statistic 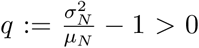 and recalling that 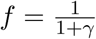 for *β* = 1:

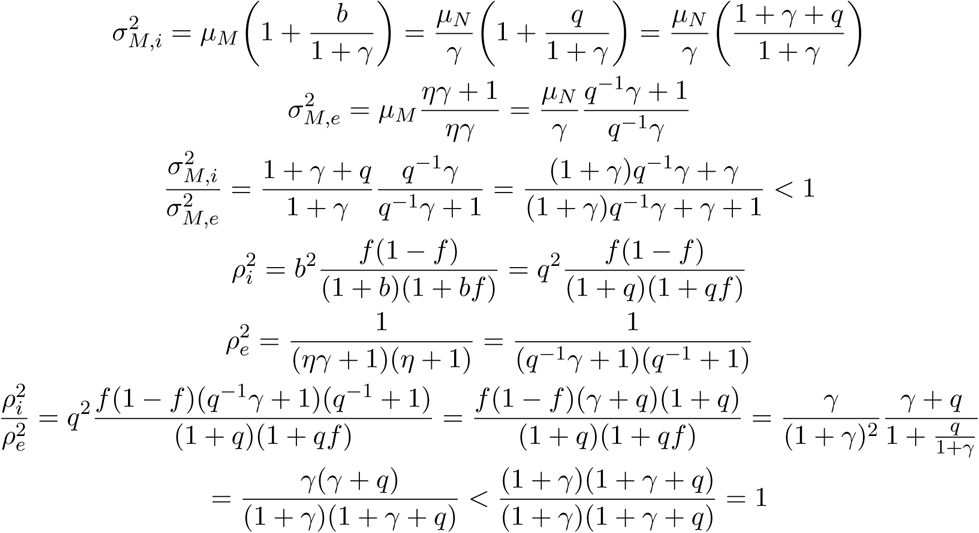

Therefore, the extrinsic noise model is overdispersed with respect to the intrinsic noise model, but its nascent and mature copy numbers are more highly correlated.

### 5.2 Case of constrained 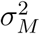

Using these expressions, it is straightforward to extend the analysis to the scenario of fixing 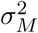:

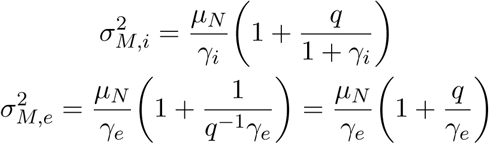

The physical solutions for *γ*_*i*_ and *γ*_*e*_ are given by positive roots of quadratic equations. For the intrinsic noise model:

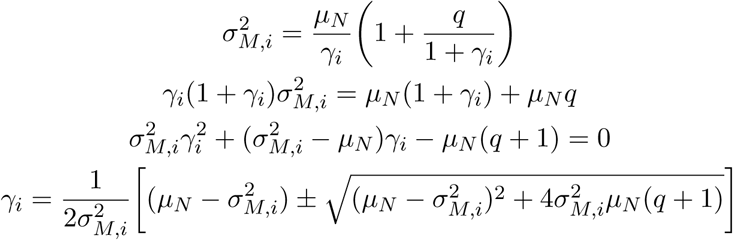

Since 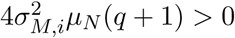, the physical solution (*γ*_*i*_ *>* 0) is given by:

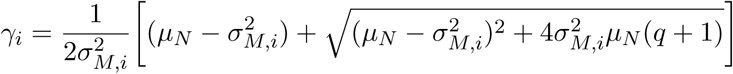

Further, for the extrinsic noise model:

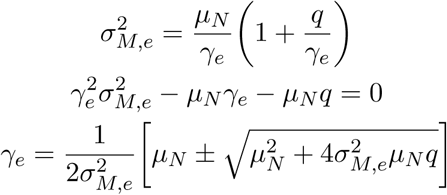

Again, since 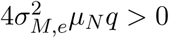, the physical solution (*γ*_*e*_ *>* 0) is given by:

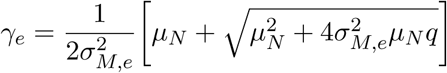

Imposing equal variances:

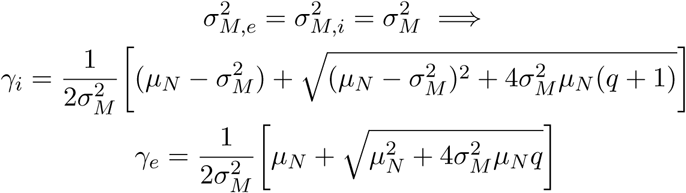

Consider 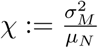. This definition yields:

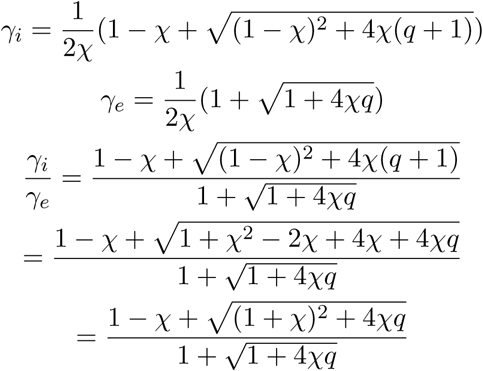

We may investigate the case where this quantity is equal to 1:

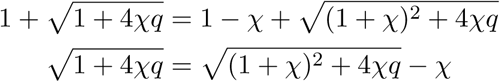

No values of *q, 𝒳 >* 0 yield this equality. This is straightforward because even the more general equation 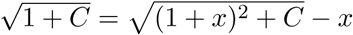 is nowhere satisfied for *x, C >* 0. Therefore, 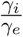 is never 1. From the quadratic equation solution, we know that *γ*_*e*_ and *γ*_*i*_ are both constrained to be positive; therefore, 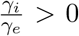. Using the test case *𝒳* = *q* = 1, we yield 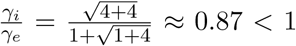. Since *γ*_*e*_ is nonzero and *γ*_*i*_(*𝒳, q*) is continuous with respect to both variables, 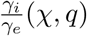 is continuous. Finally, we conclude that 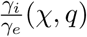 is always constrained to (0, 1) and *γ*_*e*_ *> γ*_*i*_ whenever 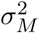 is fixed. This result matches the intuition of the provided by the finding that 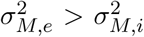 whenever *γ* is fixed: to compensate for increased dispersion in the extrinsic noise model, the degradation rate must be increased. It trivially follows that *µ*_*M,i*_ *> µ*_*M,e*_.

## 6 Experimental opportunities and limitations

Multiple experimental approaches are available for the collection of nascent and mature mRNA data. We focus on the most prevalent technologies and their relevance to the modeling question at hand.

Fluorescence microscopy methods are broadly divided between spatial transcriptomics and intron counting. Spatial transcriptomics leverages relative positions of fluorescently-labeled mRNA and DNA to identify DNA-localized nascent mRNA [22, 29]. Intron counting directly detects intron-targeted fluorescent probes [23]. These methods are rather complex and impractical to perform on a genome-wide scale. Furthermore, we are unaware of any studies combining them with dual-reporter assays to directly estimate intrinsic and extrinsic noise. Finally, the discrimination of nascent and mature mRNA aside, dual-reporter assays are in general impractical to scale to large numbers of genes.

Sequencing methods are are broadly divided between labeling and bioinformatics. Labeling refers to spiking the live media with a nucleoside analogue and distinguishing older and newer mRNA molecules based on characteristic mutations [30–33]. Purely computational methods do not require labeling, but identify nascent mRNA based on intron-aligned reads [24, 34]. These methods yield genome-wide information; however, they are not amenable to reporter duplication on the same scale. Commercially-available short-read methods present the problem of isoform indistinguishability if introns interest are outside the read region [24]. Finally, both short- and long-read methods tend to rely on the capture of polyadenylated tails [7, 35, 36], which are not present in nascent mRNA, introducing the potential of technical bias against the nascent molecules of interest. Off-target priming at intronic polyadenine sites [24, 37] and experimental methods including poly(A) ligation [31] facilitate the capture and identification of nascent transcripts, but the magnitude of technical biases is as of yet uncharacterized.

Parenthetically, we note that the motivating study by Ham et al. [13] describes a purely data-based approach to the identification of extrinsic effects, based upon the identification of heavy distribution tails. This approach appears to be quite powerful based on the provided demonstration. However, certain aspects are potentially problematic. The validation compares the tail behavior of the telegraph model to the compound telegraph model. However, even relatively simple telegraph models suffer from parameter non-identifiability issues [38, 39], so the robustness of the method is unclear. The specific fit method and metric are not reported; it is not clear that the conventional choices are appropriate when tail behavior is significant. Recent work in extreme value theory proposes several Rényi divergence alternatives. [40]. Finally, we note that the underlying data is from Zheng et al. [7], which is the earliest version of the 10X Genomics single-cell RNA sequencing platform. Since the underlying mammalian physiology has export and splicing processes [41], but 10X sequencing explicitly focuses on exonic reads [7], it is unclear that the choice of a one-stage model is justified. More problematically, raw count data are rarely used in scRNA-seq analyses [42], with substantial debate and disagreement regarding the appropriate approach to normalization [43–46]. Therefore, it is conceivable that technical biases may, in part, explain the 15-25 cells with extremely high expression that control the kernel density in the tail region, used to support the hypothesis of extrinsic noise.

## 7 Discussion

In spite of the indistinguishability of negative binomial distributions produced by intrinsic and extrinsic noise, the behavior of downstream processed gene products is substantially divergent. Specifically, we report inequality between the two models’ lower moments. Even given identical nascent marginals, it is impossible to produce identical mature marginals, and by extension full joint distributions, using the two models. More dramatically, it is impossible for the solutions’ mature marginals even to share more than one low-order moment.

In practice, if experimental joint or marginal copy-number distributions are available, it is possible to use relative likelihood testing to choose the better-fitting model. The relevant test statistic is 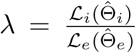, where 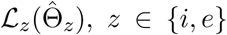 is the value of the likelihood function of the intrinsic or extrinsic model at the maximum likelihood joint parameter estimate. No closed-form joint maximum likelihood estimators are available for either model; however, estimation by numerical optimization is straightforward, especially starting at the moment-based estimates reported above. The two models’ qualitative behaviors are illustrated in Figure 2. We use the Gillespie algorithm [47] to simulate both systems given identical nascent distributions (*r* = 1.8, 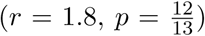) and downstream processing rates (*β* = 0.5, *γ* = 0.4). The solutions are dramatically different. As expected from the analytical moments, the extrinsic noise model gives a much more correlated joint distribution. However, despite identical marginal mature expectations, the extrinsic model has a longer tail, yielding a higher variance for that species. This drastic disagreement between distributions confirms that multimodal data is sufficient to distinguish between the two hypothesized sources of stochasticity. In addition to the theoretical and qualitative results, we provide simulation routines for both noise models. Furthermore, to facilitate comparison with discrete copy-number data, we report analytical marginal and joint distributions implied by the formulation of the system with extrinsic noise; their agreement with the simulation is shown in Figure 2. These distributions, along with moment-based initial parameter estimates, can be directly used for inference and hypothesis testing against other models.

**Figure 2:**
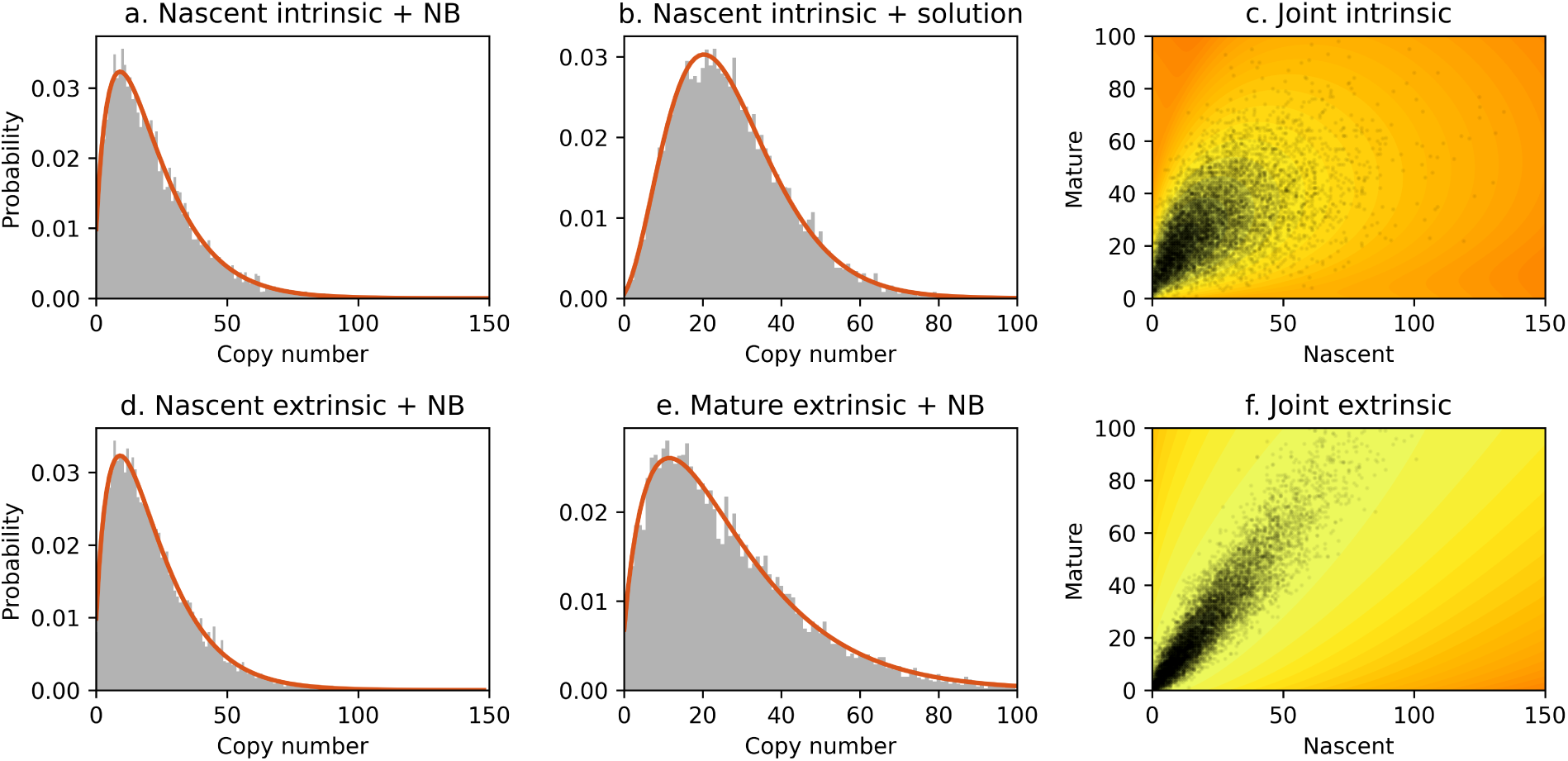
(a-c) Distributions for the intrinsic noise model with *b* = 12, *k*_*i*_ = 0.9, *β* = 0.5, and *γ* = 0.4, generated using 10^4^ simulations. (a) Nascent marginal (gray region: copy-number histogram; orange line: analytical solution). (b) Mature marginal (gray region: copy-number histogram; orange line: analytical solution). (c) Joint distribution (points: cells; color: log_10_ analytical solution, lighter color corresponds to higher probability mass). (d-f) Distributions for the extrinsic noise model with *α* = *k*_*i*_*β*^*−*1^, *η* = (*bβ*)^*−*1^, *β* = 0.5, and *γ* = 0.4, generated using 10^4^ simulations. (d) Nascent marginal (gray region: copy-number histogram; orange line: analytical solution). (e) Mature marginal (gray region: copy-number histogram; orange line: analytical solution). (f) Joint distribution (points: cells; color: log_10_ analytical solution, lighter color corresponds to higher probability mass).

Given the modeling-based insight into model identifiability, we suggest that multimodal data collection presents a valuable route to the identification of noise models. Specifically, we anticipate increased relevance for single-cell RNA sequencing, which has been challenging to integrate with experimental controls for the noise sources. Therefore, we suggest that experimental improvements in the detection of the nascent transcriptome, as well as theoretical improvements in the modeling of technical noise, would allow identification of sources of biological stochasticity on a genome-wide scale. Finally, the discrete modeling framework we discuss is immediately interpretable in terms of biophysical parameters.

## 8 Code Availability

MATLAB and Python code that can be used to reproduce Figure 2, including the simulation and plotting routines, is available at https://github.com/pachterlab/GP_2020_2.

## 9 Acknowledgments

The DNA, pre-mRNA, and mature mRNA illustrations used in Figure 1, modified from [26], are derivatives of the DNA Twemoji by Twitter, Inc., used under CC-BY 4.0. G.G. and L.P. are partially funded by NIH U19MH114830.

